# On the presence of elusive sea slugs and snails in the western Mediterranean (Mollusca: Gastropoda): new records

**DOI:** 10.1101/2020.07.29.226423

**Authors:** Xavier Salvador, Robert Fernández-Vilert, Juan Moles

## Abstract

Citizen science provides us with much information about charismatic taxa such as the opisthobranchs, thus contributing enormously to enlarging the geographic distribution of species. This study collects new records of elusive sea slugs and snails in the Western Mediterranean coast (especially in the Catalan coast and the French Mediterranean coast) and contributes to new ecological information regarding phenology, diet, and behaviour. Out of the 36 species reported here, 20 correspond to new records in the Catalan coast (NE Spain), three are new records of pelagic pteropods for the Spanish coast, and 10 other species are new records for the French Mediterranean coast. All records have been registered at the online database of the NGO named Catalan Opisthobranch Research Group (GROC). This study highlights the importance of sampling at night and, especially, in shallow, often-understudied waters, which usually gather high species diversity. We believe the high-quality pictures and related species’ information will serve future researchers and divers find and recognize these species in the Mediterranean basin.

## Introduction

Marine heterobranch gastropods, commonly named opisthobranchs, have a worldwide approximate diversity of over 6,000 species (Carefoot, 1987; Wägele *et al*., 2008). They are especially diverse in tropical and temperate waters where there is usually high mollusc diversity (Valentine & Jablonski, 2015). In the Spanish coast, 523 species have been recorded so far (Cervera *et al*., 2004) and about a half were found in the NE of the Iberian Peninsula (Catalonia), concretely 205 species (Ballesteros, *et al*., 2016). Current advances in species molecular assessments in the Mediterranean are helping to discover many new species that, in some cases, respond to hidden or cryptic speciation in taxa with similar morphological characters and, thus, difficult to discern underwater. Recent examples of such studies deal with the genus *Trinchesia* (Korshunova *et al*., 2019) for which four out of one species have been now raised; also, new species of the elusive genus *Runcina* (Araujo *et al*., 2019) or the systematic replacement of *Eubranchus farrani* (Alder & Hancock, 1844), now *Amphorina farrani*, with an additional new species described (Korshunova *et al*., 2020). Not only new species are described, also historically synonymized taxa are being resurrected to gather the new molecular diversity unveiled, such is the case of the umbraculid genus *Tylodina*, for which recent morpho-anatomical and molecular evidence suggested the resurrection of *T. rafinesquii* Philippi, 1836 (Fernández-Vilert *et al., in press*). Overall, many systematic reassessments aided by molecular tools keep revealing new taxa and further taxonomic readjustments.

During the past decades, numerous cases of allochthonous species arrivals have been detected in the Mediterranean. For instance, the aeolid *Godiva quadricolor* (Barnard, 1927) is an example of how allochthonous species are widening their range through the establishment in new areas (Cervera *et al*., 2010). Other common sightings from usually altered places with human impact are the sea hares *Bursatella leachii* Blainville, 1817 firstly reported in Israel (O’Donoghue & White, 1940) and now found all over the Mediterranean (Ibáñez-Yuste *et al*., 2012), and *Aplysia dactylomela* Rang, 1828 with an Atlantic origin (Valdés *et al*., 2013) currently found from side to side of the Mediterranean (Moles *et al*., 2017).

Because of all abovementioned reasons, heterobranch diversity and systematics in the Mediterranean are in the curve of change in the decades to come. In such a dynamic system we encourage the use of citizen science platforms to track down rare or new species (e.g., Trainito *et al*., 2017; Fernández-Vilert *et al., in press*), distribution patterns (see GROC, 2009–2020), and the spread of alien species (e.g., Fernández-Vilert *et al*., 2018). In that sense, the objective of the present study is to highlight the first or scarce new records of Heterobranchia species along the Mediterranean Spanish and French coasts using data obtained through the Catalan Opisthobranch Research Group (GROC) based on citizen science. Remarkably, the observations reported here were mainly done at shallow waters during night-time prospecting by freediving and across seasons.

## Material and Methods

Most specimens were found while freediving from 0 to 6 m depth and photographed underwater with a Nikon D90 and Nikon D7200 coupled with a 60 mm and 105 mm macro lens by X. Salvador. A map of the sampling area was generated at https://www.simplemappr.net/#tabs=0 and encompass the Southern Mediterranean coast of France (étang de Thau), the Catalan coast (NE Spain), and Granada in the South Eastern Spanish coast (Fig. 1). Species external descriptions are based on our photographic material, total length (*L*) for the analysed specimens was measured *in situ* with the aid of a ruler or in the lab under a stereomicroscope. All records were deposited in the online database of the Catalan Opisthobranch Research Group (GROC, 2009–2020).

**Figure 1.**
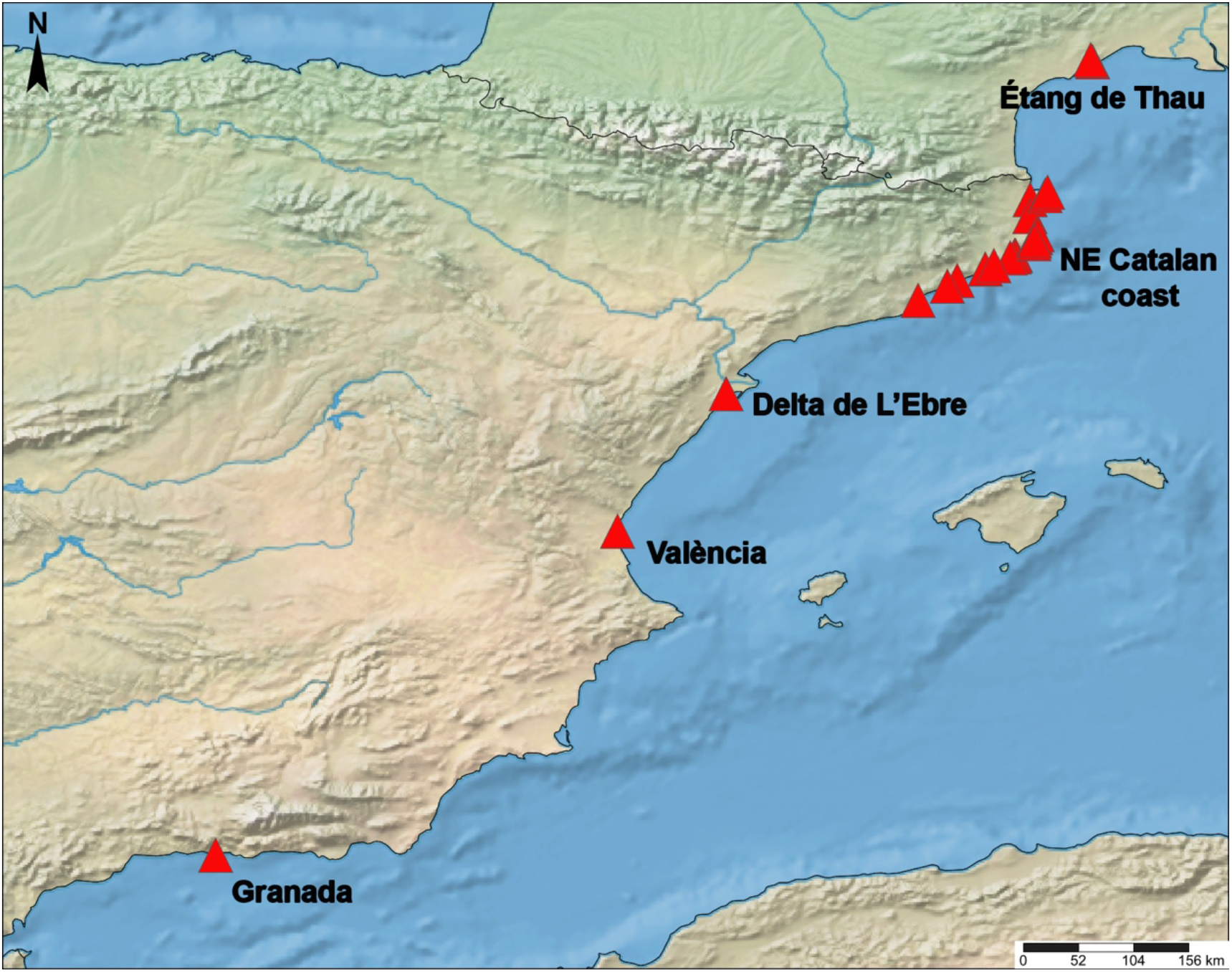
Distribution map from the Western Mediterranean coast showing the main sampling stations.

## Results

> SYSTEMATICS
>
> Order CEPHALASPIDEA P. Fischer, 1883
>
> Family HAMINOEIDAE Pilsbry, 1895
>
> Genus *Haloa* Pilsbry, 1921
>
> ***Haloa japonica* (Pilsbry, 1895)**
>
> (Figure 2A)

**Figure 2.**
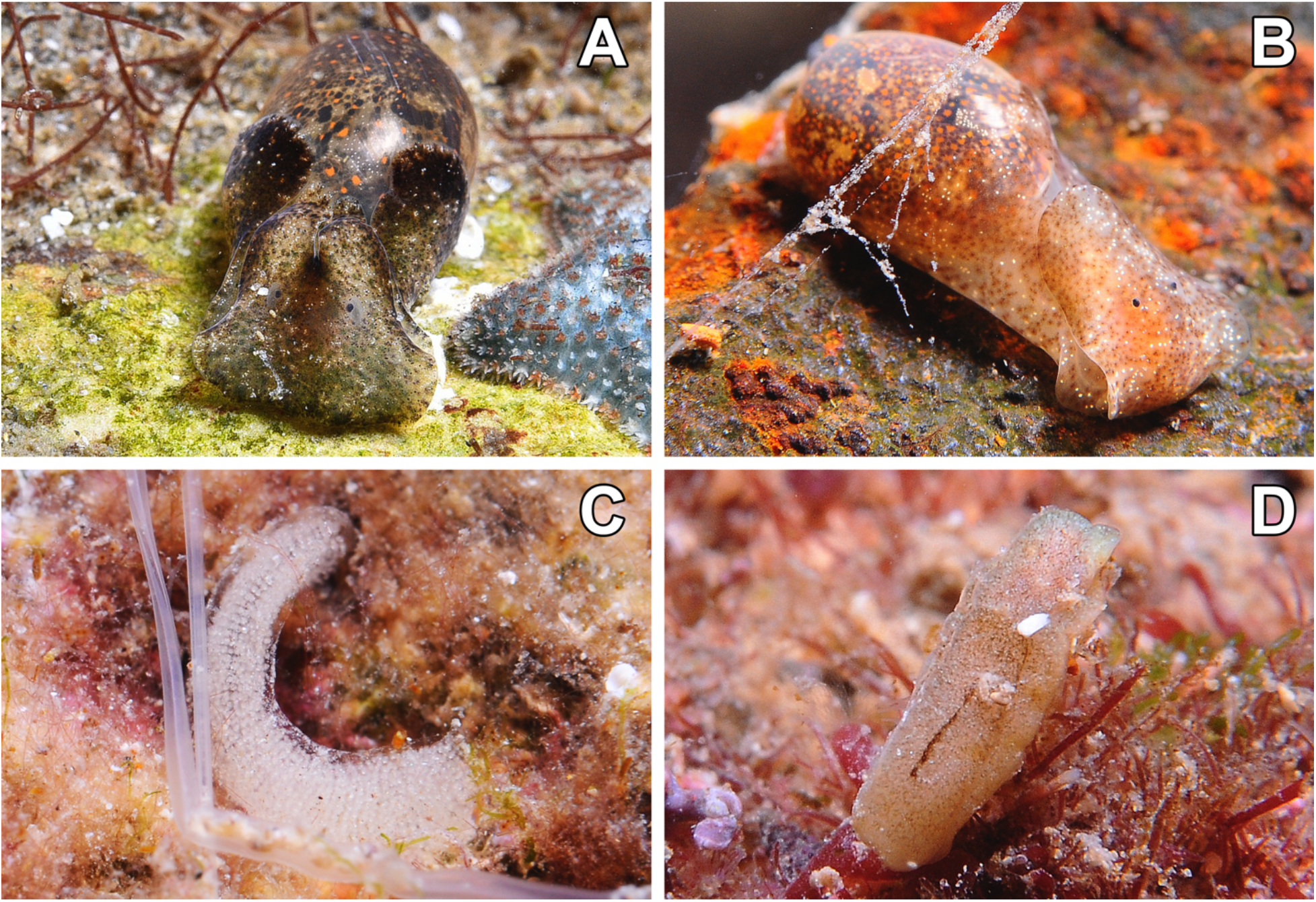
Live pictures of the Cephalaspidea species: A – *Haloa japonica*. B – *Haminoea orteai*. C – Egg mass of *Haminoea orteai*. D – *Philine catena*.

### Material examined

Le Ponton, étang de Thau, Sète (France), 43°25’28.5”N, 3°42’E, 3-Nov-2017, 2.1 m depth, 1 spc., adult, *L* = 20 mm; 14-Apr-2018, 1.7 m depth, 3 spcs, adults, *L* = 20–30 mm; 21-May-2018, 1.9 m depth, 8 spcs, adults, *L* = up to 35 mm.

### External morphology

Shell external, translucent, background colour brown with big orange punctuations. Parapodia darker on top. Head lighter on edge, with darker central part between the eyes.

### Ecology

Found at night on soft bottoms, between Ceramiales species and other undetermined red algae. The groups of specimens found during April and May were mating.

### Distribution

Japanese coast (Pilsbry, 1895); USA (GBIF, 2020); Spain: Galicia (Cervera *et al*., 2004), Cantabria (Cervera *et al*., 2004), Andalucía (Cervera *et al*., 2004); France (this study).

### Remarks

In the area there are species of Japanese origin, such as the alga Wakame *[Undaria pinnatifida* (Harvey) Suringar, 1873] or the oyster [*Magallana gigas* (Thunberg, 1793)], so their presence may be justified by aquaculture.

> ***Haminoea orteai* Talavera, Murillo & Templado, 1987**
>
> (Figure 2B,C)

### Material examined

Cala Maset caves, Sant Feliu de Guíxols (Spain), 41°47’10.5”N, 3°2’44.6”E, 16-Oct-2017, 0.6 m depth, >40 spcs, juveniles, adults, and egg masses, *L* = 2–15 mm; 31-Dec-2017, 1.2 m depth, 3 spcs, juveniles and adults, *L* = up to 15 mm; Le Ponton, étang de Thau, Sète (France), 43°25’28.5”N, 3°42’E, 3-Nov-2017, 1.4 m depth, 2 spcs, adults, *L* = 20 mm; Fòrum pools, Barcelona (Spain), 41°24’34.5”N, 2°13’36.8”E, 25-Nov-2017, 1.7 m depth, 2 spcs, adults and egg masses, *L* = 20 mm; 27-Apr-2018, 0.8 m depth, 1 spc., adult, *L* = 10 mm; l’Espigó, Roses (Spain), 42°15’39.8”N, 3°10’26.8”E, 28-Feb-2020, 0.8 m depth, 5 spcs, adults and egg masses, *L* = 25 mm.

### External morphology

Shell external, translucent, background colour light brown to grey with dark brown, white and orange punctuations surrounded of little dark red points. Parapodia lighter in colour. Head lighter in colour, area around eyes pigmented.

### Ecology

Specimens were found crawling on calcareous rocks with undetermined green algae. Egg masses were observed in November and juveniles in December. Egg masses elongated, forming a “C”, with white eggs (Fig. 1C).

### Distribution

Açores (Cervera *et al*., 2004); Spain: Canary Islands (Cervera *et al*., 2004), Andalucía (Cervera *et al*., 2004), Spanish levant (Cervera *et al*., 2004; Talavera *et al*., 1987), Catalonia (this study); France (this study).

### Remarks

*Haminoea* species in the Mediterranean are yet poorly studied (M. A. E. Malaquias pers. comm.) and thus, the morphological characters to distinguish the species are not clear. The visual distinguishing feature of this species is the presence of a pigmented periocular area. The presence of small orange dots under the shell surrounded by smaller dots of dark purple or black colour, forming a flower pattern is a pattern of colouration that found in all studied specimens.

> Family PHILINIDAE Gray, 1850 (1815)
>
> Genus *Philine* Ascanius, 1772
>
> ***Philine catena* (Montagu, 1803)**
>
> (Figure 2D)

### Material examined

Cala Maset caves, Sant Feliu de Guíxols (Spain), 41°47’10”N, 3°2’44”E, 13-Dec-2017, 1.3 m depth, 1 spc., adults, *L* = 5 mm; Punta del Romaní, L’Escala (Spain), 42°6’54”N, 3°10’9”E, 30-Dec-2017, 2.5 m depth, 2 spcs, adults, *L* = 3–7 mm; Cala d’Aiguafreda, Begur (Spain), 41°57’49”N, 3°13’41”E, 31-Jan-2018, 1.2 m depth, 1 spc., adult, *L* = 5 mm.

### External morphology

Body elongate, narrow, white-beige or brown in colour, with a discontinuous dark-brown band over cephalic shield. Cephalic shield longer than shell. Parapodia short, not overlapping. Shell completely covered by mantle. Probably individuals are hiding among the sediment grains during the day and actively moving at night.

### Ecology

The specimens were found crawling at night on rocks with algae and evident sediment loading between rocky outcrops.

### Distribution

North Sea (GBIF, 2020); Açores (Cervera *et al*., 2004); Portugal (Cervera *et al*., 2004); Spain: Galicia (Cervera *et al*., 2004), Cantabria (Cervera *et al*., 2004), Catalonia (this study).

### Remarks

Molecular and morphological data reveals this species may not belong to the genus *Philine* (J. Moles unpubl. data).

> Order NUDIBRANCHIA Cuvier, 1817
>
> Family AEGIRIDAE P. Fischer, 1883
>
> Genus *Aegires* Lovén, 1844
>
> ***Aegires palensis* Ortea, Luque & Templado, 1990**
>
> (Figure 3A)

**Figure 3.**
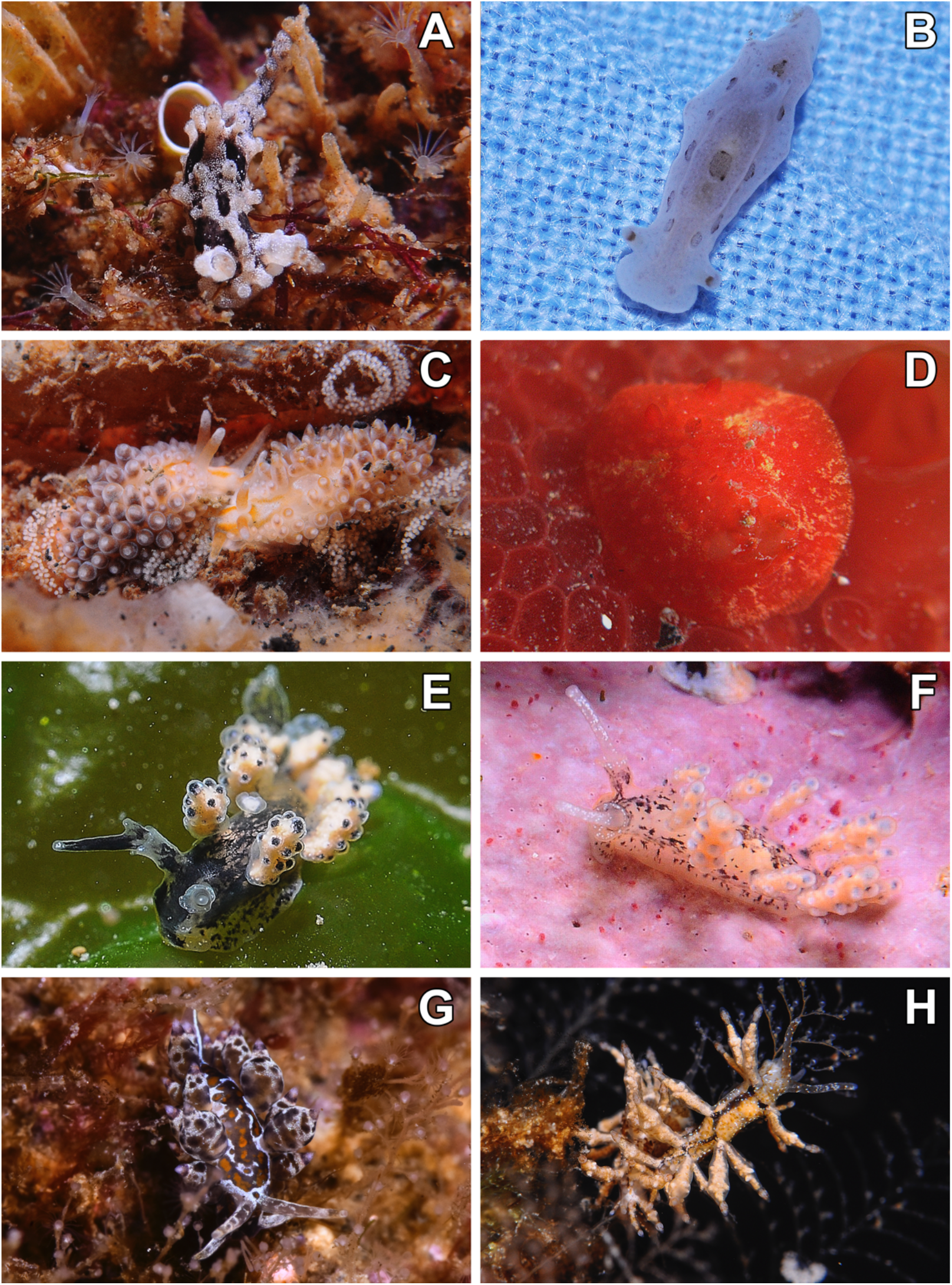
Live pictures of the Nudibranchia species: A – *Aegires palensis*. B – *Aegires sublaevis*. C – Two specimens of *Anteaeolidiella lurana* and their egg masses. D – *Aldisa smaragdina* on top of the sponge *Phorbas topsenti*. E – *Doto cervicenigra*. F – *Doto pygmaea*. G – *Amphorina andra* surrounded by hydrozoans of the genus *Sertularella*. H – *Eubranchus prietoi* on the hydrozoan *Kirchenpaueria halecioides*.

### Material examined

Cala Ventosa, Sant Feliu de Guíxols (Spain), 41°47’5”N, 3°2’52”E, 13-Sep-2017, 1 m depth, 1 spc., adult, *L* = 10 mm; Cala Maset caves, Sant Feliu de Guíxols (Spain), 41°47’10”N, 3°2’44”E, 7-May-2018, 1 m depth, 7spcs, juveniles and adults, *L* = 3–15 mm; Cala d’Aiguafreda, Begur (Spain), 41°57’49”N, 3°13’41”E, 25-May-2018, 1 m depth, 6spcs, juveniles and adults, *L* = 6–15 mm.

### External morphology

Body elongate, rough, angulated, with tubercles in the laterals; body colour beige, with small white and light–brown dots; rhinophores, rhinophoral sheaths and apical part of tubercles with dark-brown spots. Branchial leaves protected by three tubercles, curved internally, equal in length.

### Ecology

All specimens were found on top of white calcareous sponges of the genus *Sycon* (pss. *S. elegans* (Bowerbank, 1845) and *S. raphanus* Schmidt, 1862) and *Ascandra contorta* (Bowerbank, 1866). Generally found at night at the entrance or inside caves, crawling on walls.

### Distribution

Spain: Andalucía, Spanish levant (Cervera *et al*., 2004), and Catalonia (Cervera *et al*., 2004; Ballesteros *et al*., 2012–2020; this study).

### Remarks

This species can be differentiated from the other Atlantic species of *Aegires* in the structure and disposition of the tubercles, body colour, and the dark brown dots in the sheaths and apex of the rhinophores (Ortea *et al*., 1990).

> ***Aegires sublaevis* Odhner, 1932**
>
> (Figure 3B)

### Material examined

Cala Maset caves, Sant Feliu de Guíxols (Spain), 41°47’10”N, 3°2’44”E, 16-Feb-2018, 1 m depth, 1 spec, adult, *L* = 15 mm.

### External morphology

Body elongate, rough, angulated with central part of dorsum flattened, with black marks in dorsum and laterals; body colour white yellowish. Branchial leaves protected by three tubercles, the central one bigger than the laterals. Rhinophores smooth, presenting a brown ring close to the top.

### Ecology

Our specimen was found feeding on white calcareous sponges of the genus *Ascandra* and, thus, displaying a whiter colouration.

### Distribution

Madeira (Malaquias *et al*., 2001); Açores (Calado, 2002; Fahey & Gosliner, 2004); Spain: Canary Islands (Odhner, 1931; Altimira & Ros, 1979, as *Serigea sublaevis*; Pérez Sánchez, Ortea, & Bacallado, 1990; Ortea, et al., 1996; 2001; 2003; Malaquias & Calado, 1997), Spanish Levant (Templado *et al*., 1987), and Catalonia (this study).

### Remarks

According to Ortea *et al*. (1996) its colouration can vary from yellow, beige or white depending on the diet, generally found feeding on *Clathrina coriacea* (Montagu, 1814).

> Family AEOLIDIIDAE Gray, 1827
>
> Genus *Anteaeolidiella* M. C. Miller, 2001
>
> ***Anteaeolidiella lurana* (Ev. Marcus & Er. Marcus, 1967)**
>
> (Figure 3C)

### Material examined

Badia dels Alfacs port, Sant Carles de la Ràpita, Montsià (Spain), 40°37’1”N, 0°35’52”E, 1.3 m depth, 3 spcs, adults, and egg masses, *L* = 15 mm; Valencia’s port (Spain), 39°27’39.9”N, 0°19’29.5”W, 18-Jan-2020, 0.2 m depth, 1 spec, adult, H= 10mm.

### External morphology

Body elongate, narrow, with a white mark surrounded by orange in dorsum; background colour white, with 2 orange lines in head, united behind eyes. Cerata short, round, background colour grey, white on top. Rhinophores long, equal, white in colour with orange central region.

### Ecology

The specimens found in Alfacs bay (Catalonia) were found under rocks mating and laying eggs. However, the specimen found in Valencia was on a vertical wall in the port, moving around hydrozoans at night, thus the species likely presents nocturnal activity.

### Distribution

Brazil (Marcus & Marcus, 1967); Italy (Ballesteros *et al*., 2012–2020); Spain: Balearic Islands (GROC, 2009–2020), Catalonia (this study).

### Remarks

The species was described from Brazil and records of its first invasion in the Mediterranean correspond to Italy by Alberto Piras (Cagliari, Italy (1983); Köhler, 2020). The two locations where it was found in this study have fairly eutrophic waters and ship activity, which may be related to its appearance.

> Family CADLINIDAE Bergh, 1891
>
> Genus *Aldisa* Bergh, 1878
>
> ***Aldisa smaragdina* Ortea, Pérez & Llera, 1982**
>
> (Figure 3D)

### Material examined

Cala d’Aiguafreda, Begur (Spain), 41°57’49”N, 3°13’41”E, 24-Apr-2015, 3 m depth, 1 spc., *L* = 7 mm.

### External morphology

Body rounded, flat; light red in colour, with two dark circles on dorsum; yellow-brown line present at laterals of body next to the first circle. Gills light red, white on top of branchial leaves.

### Ecology

The specimen was found at a shallow depth under a rock, on top of the sponge *Phorbas topsenti* Vacelet & Pérez, 2008.

### Distribution

Spain: Canary Islands (Pruvot-fol (1953) as *Aldisa binotata*; Ortea *et al*., 1982), Catalonia (this study).

### Remarks

In its original description the body was described as red with 2 big dark circles and a white line just after the first dark circle. The specimen studied here presents a lighter red colouration and lacks the second dark circle. The gills are also light red and white on the top, while the conspecific *A. banyulensis* (Pruvot-Fol, 1951) presents dark red gills, lacking the white top. Moreover, our specimen presents a yellow line in the anterior part of the body, just after the first circle and diagonally towards the front, while in *A. banyulensis* the yellow line is found between the first and second dots, displayed slightly towards the posterior part.

> Family DOTIDAE Gray, 1853
>
> Genus *Doto* Oken, 1815
>
> ***Doto cervicenigra* Ortea & Bouchet, 1989**
>
> (Figure 3E)

### Material examined

Cala Maset caves, Sant Feliu de Guíxols (Spain), 41°47’10.5”N, 3°2’44.6”E, 13-Apr-2016, 0.3 m depth, 1 spc., *L* = 5 mm; 31-Dec-2017, 1.2 m depth, 3 spcs, juveniles and adults, *L* = 15 mm; 13-Jan-2018, 0.4 m depth, 2 spcs, *L* = 3–7 mm; 31-Dec-2017, 1.2 m depth, 3 spcs, juveniles and adults, *L* = 15 mm; Le Ponton, étang de Thau, Sète (France), 43°25’28.5”N, 3°42’E, 13-Apr-2018, 0.2 m depth, 4 spcs, adults, *L* = 7–10 mm; 21-May-2018, 0.2 m depth, 12 spcs, adults and egg masses, *L* = 7–10 mm; la Farge, étang de Thau, Sète (France), 43°25’48”N, 3°42’14”E, 21-May-2018, 0.2 m depth, 17 spcs, adults and egg masses, *L* = 7-10 mm; Port de Sant Feliu (Spain), 41°46’42.2”N, 3°02’16.4”E, 29-Jan-2020, 0.2 m depth, 11 spcs, adults, juveniles, and egg masses, *L* = 2–12 mm; Arenys de Mar port (Spain), 41°34’38.8”N, 2°33’23.5”E, 15-Feb-2020, 0.2 m depth, 2 spcs, adults, *L* = 10 mm; Port de Blanes (Spain), 41°40’25.5”N, 2°47’48.6”E, 2-Mar-2020, 0.2 m depth, 2 spcs, adults, *L* = 6–10 mm.

### External morphology

Body elongate, narrow, background colour white with black marks all along but more concentrated in the head area. Rhinophores black with white tips. Cerata with tubercles, apical part black.

### Ecology

The specimens were found at shallow depths, influenced by fresh water, on top of rocks and hydrozoans, possibly *Obelia* sp., where they were feeding and laying the eggs (short, lineal, with big white eggs) at the bottom of the colony. They were found more active during the night.

### Distribution

Corsica (Ortea & Bouchet, 1989); Italy (Ballesteros *et al*., 2012–2020); Spain: Mallorca (GROC, 2009–2020), Catalonia (this study); France (this study).

### Remarks

This species is easily distinguished from other conspecific *Doto* species by the presence of black rhinophores and the small tubercles with a black dot in the cerata (Ortea & Bouchet, 1989).

> ***Doto pygmaea* Bergh, 1871**
>
> (Figure 3F)

### Material examined

Cala d’Aiguablava, Begur (Spain), 41°56’13.4”N, 3°13’08.6”E,22-Jun-2018, 0 m depth, 6 spcs, juvenils and adults, *L* = 2–5 mm; Cala Maset caves, Sant Feliu de Guíxols (Spain), 41°47’10.5”N, 3°2’44.6”E, 11-Feb-2020, 0.1 m depth, 7spcs, adults and egg masses, *L* = 5–10 mm.

### External morphology

Body elongate, narrow, background colour white with black patches to completely black. Rhinophores elongate, translucid white. Cerata displayed in a sinuous “S” fashion with white tubercles only in external part, the inner part being almost smooth.

### Ecology

All specimens were found over floating debris. The specimens from Aiguablava were living over a plastic cube eating hydrozoan colonies, probably of the genus *Obelia*. The specimens of Cala Maset were living in a floating wood that appeared after a storm. They were mating and laying egg masses, while eating hydrozoans of the genus *Tubularia*. Their egg masses are shaped like a winding cord forming an “S”, the eggs are yellowish.

### Distribution

Italy (Schmekel & Portmann, 1982); Spain: Canary Islands (Cervera *et al*., 2004), Levantine Spanish coast (Cervera *et al*., 2004), Catalonia (this study).

### Remarks

This species can be differentiated from other *Doto* species by having a black body, with very elongated rhinophores, a smooth edge on the rhinophoral sheath, and “S” shaped cerata, the inner part being almost smooth and the external with tubercles aligned longitudinally (Ortea *et al*., 1997).

> Family EUBRANCHIDAE Odhner, 1934
>
> Genus *Amphorina* Quatrefages, 1844
>
> ***Amphorina andra* Korshunova, Malmberg, Prkić, Petani, Fletcher, Lundin, Martynov, 2020**
>
> (Figure 3G)

### Material examined

Cala d’Aiguafreda, Begur (Spain), 41°57’49”N, 3°13’41”E, 22-Jun-2013, 18 m depth, 1 spec, juvenile, *L* = 3 mm (GROC 2013); Balaruc-les-Bains (France), 43°26’28.8”N, 3°41’3”E, 5-Apr-2014, 2 m depth, 2 spcs, adults, *L* = 10–15 mm; Barra de l’Arbre, Mataró (Spain), 41°31’57.9”N, 2°28’21.2”E, 1-May-2014, 20 m depth, 1 spec, juvenile, H= 2 mm; Punta del Romaní, L’Escala (Spain), 42°6’54”N, 3°10’9”E, 12-Apr-2015, 4 m depth, 3 spcs, adults, *L* = 10–12 mm; Cala Maset caves, Sant Feliu de Guíxols (Spain), 41°47’10”N, 3°2’44”E, 16-Jan-2018, 1 m depth, 2 spcs, adults, *L* = 10 mm; 16-Jan-2018, 1 m depth, 2 spcs, adults and egg masses, *L* = 10–12 mm; 22-Jan-2018, 0.7 m depth, 6spcs, adults, mating, and egg masses, *L* = 12–15 mm; 16-Feb-2018, 1.2 m depth, 2 spcs, adults, *L* = 8–10 mm; 9-Mar-2018, 0.6 m depth, 6 spcs, adults, mating and egg masses, *L* = 10–15 mm; Punta d’en Bosch, Sant Feliu de Guíxols (Spain), 41°45’54”N, 3°0’11”E, 17-Apr-2018, 1 m depth, 5 spcs, adults mating, *L* = 8–12 mm; Cova de l’infern (Spain), 42°19’2.75”N, 3°19’12.74”E, 2-Mar-2019, 1 m depth, 3 spcs, adults and egg masses, *L* = 10–20 mm.

### External morphology

Body elongate, narrow, background colour variable, normally white with black and orange marks, some specimens displaying a completely white or orange colouration. Rhinophores and oral tentacles smooth, yellow and white apically. Cerata globular, with white and black marks, apical part yellow.

### Ecology

The specimens were found in broad range of depths, always associated to hydrozoan colonies. The specimens of Sant Feliu were found inside a cave, in presence of hydrozoans of the genus *Sertularella*, where they often lay the egg masses. They were most active overnight, mating and laying eggs. During the day they rest at the base of the hydrozoans. The egg mass is spiral shaped, usually laid on rocks, although they can also be on the hydrozoans upon which they feed.

### Distribution

Italy and Croatia (Korshunova *et al*., 2020); France (this study); Spain: Catalonia (this study).

### Remarks

This species have been recently described and was formally attributed to its conspecific and sympatric *A. farrani* (Alder & Hancock, 1845; formerly *Eubranchus farrani*). *Amphorina andra* presents a very variable colouration pattern, from completely white or orange, to a white body with black spots and orange circles on the back. Specimens found at deeper depths were typically small in size, whereas specimens found at very shallow depths, mostly in caves, were larger and found on hydrozoan colonies of the genus *Obelia* and *Sertularella*, among others. The populations observed in Croatia were very shallow, between 0 and 0.5 m (Korshunova *et al*., 2020), as was the population studied in Sant Feliu, Catalonia (Spain).

> Genus *Eubranchus* Forbes, 1838
>
> ***Eubranchus prietoi* Llera & Ortea, 1981**
>
> (Figure 3H)

### Material examined

Cala Maset caves, Sant Feliu de Guíxols (Spain), 41°47’10”N, 3°2’44”E, 11 to 16-Feb-2018, 1.5–2 m depth, 13spcs, adults and egg masses, *L* = 4–7 mm; 9 to 15-Mar-2018, 1.5 m depth, 13spcs, juveniles, adults, and egg masses, *L* = 6–10 mm; 30-Mar-2018, 1.5 m depth, 7 spcs, juveniles, adults, and egg masses, *L* = 7–15 mm; 15-Apr-2018, 1.5 m depth, 2 spcs, adults, *L* = 8 mm; 7-May-2018, 0.7 m depth, 2 spcs, adults, *L* = 7 mm.

### *External* morphology

Body elongate, narrow, with a black mark in dorsum (corresponding to digestive gland seen by transparency), starting between the first couple of cerata and finishing at the last group of cerata. Background colour beige with brown and white punctuation. Rhinophores long, equal. Cerata large and fusiform compressed at the middle, transparent with white to dark brown visible digestive gland, white tips.

### Ecology

The specimens were found during the day resting at the base of the hydrozoan *Kirchenpaueria halecioides* (Alder, 1859), while actively crawling, mating, and feeding on the colony at night. C-shaped egg masses were found over the hydrozoan.

### Distribution

Gibraltar (García-Gómez, 1987); Spain: Cantabria (Ortea & Bacallado, 1981), Catalonia (this study).

### Remarks

When active, they present the cerata completely lateralized, camouflaging themselves around the pycnogonids also found on the hydrozoan colonies (X. Salvador pers. obs.). It can be differentiated externally from other *Eubranchus* species by the insertion of the first cerata groupings and the visible black digestive gland on dorsum between cerata (Llera & Ortea, 1981).

> Family FACELINIDAE Bergh, 1889
>
> Genus *Godiva* Macnae, 1954
>
> ***Godiva quadricolor* (Barnard, 1927)**
>
> (Figure 4A)

**Figure 4.**
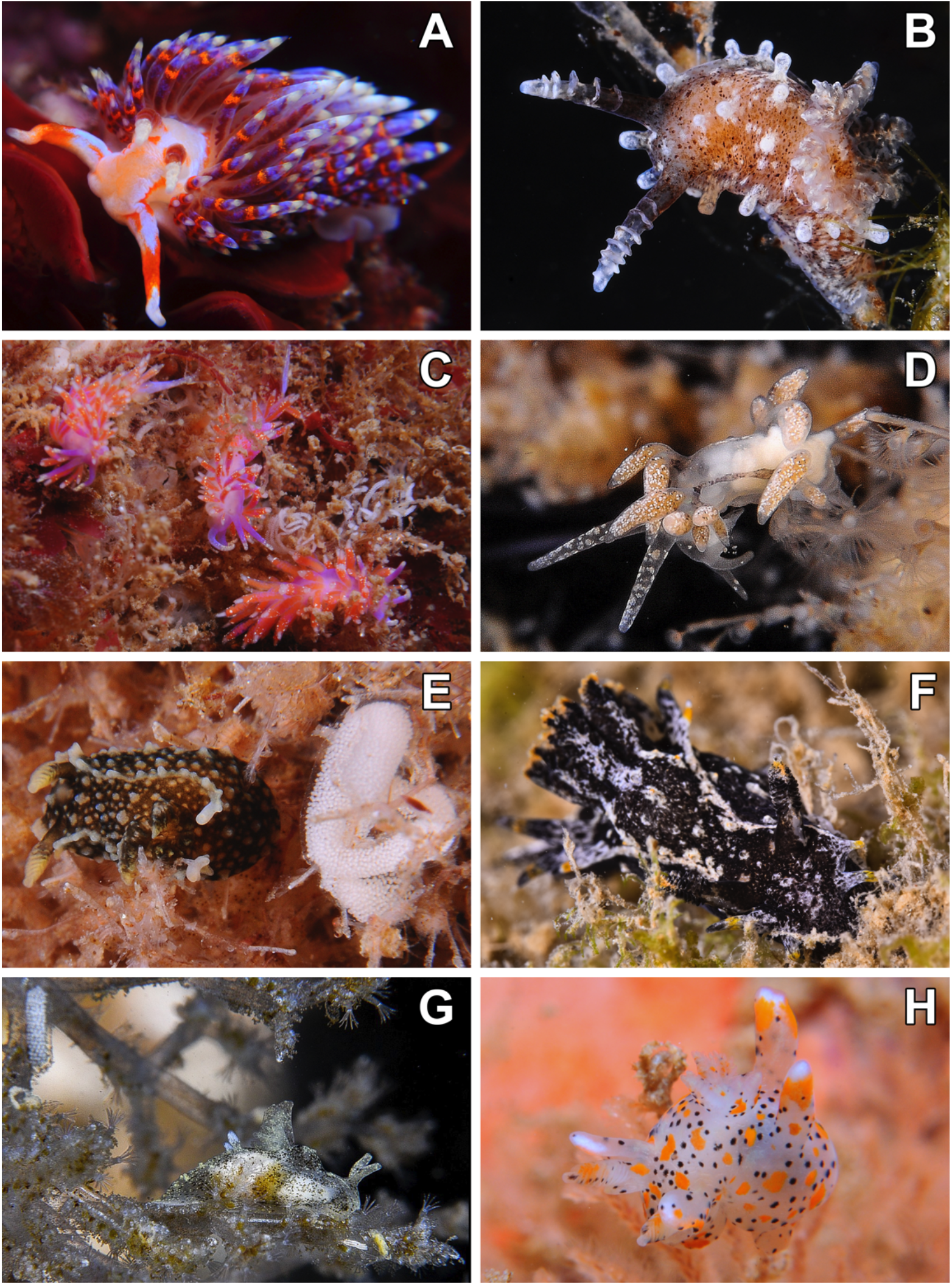
Live pictures of the Nudibranchia species: A – *Godiva quadricolor*. B – *Okenia longiductis* on the bryozoan *Amathia verticillata*. C – Three specimens of *Piseinotecus soussi* and their egg massess. D – *Piseinotecus sphaeriferus* on top of the hydrozoan *Obelia* sp. E – *Palio nothus* and its egg mass. F *-Polycera hedgpethi*. G – *Polycerella emertoni* and egg mass on *Amathia verticillata*. H – *Thecacera pennigera* on top of the bryozoan *Bugula* sp.

### Material examined

Cala Maset caves, Sant Feliu de Guíxols (Spain), 41°47’10.5”N, 3°2’44.6”E, 31-Dec-2017, 2 m depth, 1 spc., *L* = 10 mm; 5-Jan-2018, 1.3 m depth, 1 spc., adult, *L* = 25 mm; 22-Jan-2018, 1.3 m depth, 1 spc., adult, *L* = 25 mm; 29-Jan-2018, 1.3 m depth, 1 spc., adult, *L* = 25 mm.

### External morphology

Body elongate, narrow, background colour light orange with whitish blue electric marks. Oral tentacles presenting a whitish blue line connecting apical part with base of rhinophores. Rhinophores equal, white yellowish apically. Cerata abundant, smooth, coloured in red at base, orange, blue and yellow towards tip.

### Ecology

This species has a very wide diet (Betti *et al*., 2015), commonly found eating anemones (e.g. the genera *Anemonia* and *Aiptasia*), worms (e.g. *Sabella*), and other heterobranchs such as *Spurilla neapolitana* (Delle Chiaje, 1841).

### Distribution

This is an invasive species originally from South Africa and the Pacific Sea (GBIF, 2020), now found in Italy, France, Andalucía (Ballesteros *et al*., 2012–2020), and Catalonia (this study).

### Remarks

This species is a big facelinid with a very characteristic colour pattern, difficult to misidentify. In the étang de Thau the species is very abundant during Spring.

> Family GONIODORIDIDAE H. Adams & A. Adams, 1854
>
> Genus *Okenia* Menke, 1830
>
> ***Okenia longiductis* Pola, Paz-Sedano, Macali, Minchin, Marchini, Vitale, Licchelli & Crocetta, 2019**
>
> (Figure 4B)

### Material examined

Cala Maset caves, Sant Feliu de Guíxols (Spain), 41°47’10”N, 3°2’44”E, 23-Sep-2016, 1 m depth, 5 spcs, juveniles, adults, and egg masses, *L* = 2–8 mm; 14-Sep-2017, 1 m depth, 7 spcs, adults and egg masses, *L* = 5–13 mm; 16-Oct-2017, 1 m depth, > 20 spcs, adults and egg masses, *L* = 7–13 mm; Cala Ventosa, Sant Feliu de Guíxols (Spain), 41°47’5”N, 3°2’52”E, 13-Sep-2017, 1 m depth, 2 spcs, adults and egg masses, *L* = 5 mm; Fòrum pools, Barcelona (Spain), 41°24’34.5”N, 2°13’36.8”E, 23-Nov-2017, 2.3 m depth, 7 spcs, adults and egg masses, *L* = 5–10 mm.

### External morphology

Body elongate, narrow, background colour light brown or white with irregular brown, black and white punctuations in adults. Papillae smooth, found scattered on dorsum and laterals. Rhinophores presenting a smooth front surface, lamellar in back. Discoidal branchial leaves.

### Ecology

All specimens found at each locality were found on the bryozoans *Amathia verticillata* (Delle Chiaje, 1822), laying their egg masses.

### Distribution

Recently described in Italy and France (Pola *et al*., 2019); Spain: Catalonia (this study).

### Remarks

Pola *et al*. (2019) recently confirmed the differences between this species and *O. zoobotryon* (Smallwood, 1910), previously recorded in the Mediterranean but originally described from Bermudas. Molecular and morphological analyses in the previous study revealed differences in the reproductive system and the gill branches between these two species. Thus, according to Pola *et al*. (2019) previous records of *O. zoobotryon* in the Mediterranean are misidentifications of *O. longiductis*, and *O. zoobotryon* is only present in the western Atlantic.

> Family PISEINOTECIDAE Edmunds, 1970
>
> Genus *Piseinotecus* Er. Marcus, 1955
>
> ***Piseinotecus soussi* Tamsouri, Carmona, Moukrim & Cervera, 2014**
>
> (Figure 4C)

### Material examined

Cala Maset caves, Sant Feliu de Guíxols (Spain), 41°47’10”N, 3°2’44”E, 22-Jan-2018, 1.8 m depth, 7 spcs, juveniles, adults, and egg masses, *L* = 4–25 mm; 29-Jan-2018, 1.6 m depth, 9spcs, adults and egg masses, *L* = 12–25 mm.

### External morphology

Body elongate, narrow, background colour violet or pink. Rhinophores and oral tentacles smooth, white apically with a degraded white punctuation. Cerata smooth, long, translucent, with digestive gland visible in orange; white apically, presenting a profuse white punctuation.

### Ecology

Specimens were found mating and laying the egg masses on top of unidentified species of hydrozoans at night and hiding at the base of the colonies during daylight.

### Distribution

Morocco (Tamsouri *et al*., 2014); Italy (Ballesteros *et al*., 2012–2020), Catalonia (Ballesteros *et al*., 2016; this study).

### Remarks

Originally described from Morocco, it is quite widespread on the Catalan coast. Differs from other similar-looking species, such as *Edmunsella pedata* (Montagu, 1816), by the presence of abundant white spots in the cerata, rhinophores, and oral tentacles.

> ***Piseinotecus sphaeriferus* (Schmekel, 1965)**
>
> (Figure 4D)

### Material examined

Punta del Romaní, l’Escala, Girona (Catalunya), 42°6’53.088”N, 3°10’6.765”E, 8-November-2014, 1 spc.; Le Ponton, étang de Thau, Sète (France), 43°25’28.5”N, 3°42’E, 20-May-2018, 0.4 m depth, 1 spc., adult, *L* = 13 mm.

### External morphology

Body elongate, narrow, background colour transparent white. Rhinophores and oral tentacles smooth, translucent with white spots. Cerata smooth, background colour white or beige, base presenting a green iridescent reflection; first and second row of cerata connected with a dark line.

### Ecology

The singleton was found on top of the hydrozoan *Obelia* sp. during the night.

### Distribution

Ghana (Edmunds, 1977); Portugal (Cervera *et al*., 2004); Italy and Adriatic Sea (Ballesteros *et al*., 2012–2020); Spain: Canary Islands (Ortea *et al*., 2003), Catalonia (Ballesteros *et al*., 2016; GROC, 2009– 2020); France (this study).

### Remarks

Elusive species easily diagnosed by a green iridescent sphere at the base of the cerata (Ortea *et al*., 2003).

> Family POLYCERIDAE Alder & Hancock, 1845
>
> Genus *Palio* Gray, 1857
>
> ***Palio nothus* (Johnston, 1838)**
>
> (Figure 4E)

### Material examined

Cala Maset caves, Sant Feliu de Guíxols (Spain), 41°47’10”N, 3°2’44”E, 16-Jan-2018, 1.7 m depth, 5 spcs, adults and egg masses, *L* = 8–14 mm; 7-May-2018, 1.4 m depth, 1 spc., adult and egg masses, *L* = 15 mm; *L* = 8–14 mm; 11-Mar-2020, 2 m depth, 7 spcs, juveniles.

### External morphology

Body short, thick, background colour black with numerous white papillae. Mantle margin white. Rhinophores presenting lamellae, beige or white in colour. Branchial leaves dark brown.

### Ecology

The specimens were found mating and laying the egg masses on top of the bryozoans *Amathia lendigera* (Linnaeus, 1758) on walls. The egg masses are white with a soft brown tone in a “C” shape, laid over the bryozoans. Juvenile specimens only found in 2020 also on top of *A. lendigera*. They are more active overnight and go unnoticed during the day.

### Distribution

North Sea (England and Norway; Johnston, 1838); Spain: Catalonia (Ballesteros *et al*., 2016; this study).

### Remarks

This species has an amphi-Atlantic and boreo-artic distribution, but its identity may need confirmation since is easily confused with *Palio dubia* (M. Sars, 1829) and several records may have been misidentified (Thompson & Brown, 1984).

> Genus *Polycera* A. E. Verrill, 1880
>
> ***Polycera hedgpethi* Er. Marcus, 1964**
>
> (Figure 4F)

### Material examined

Le Ponton, étang de Thau, Sète (France), 43°25’28.5”N, 3°42’E, 8-Apr-2017, 1.2 m depth, 7 spc., adults and egg masses, *L* = 10–25 mm; 4-Oct-2017, 0.8 m depth, 3 spcs, adults, *L* = 20–25 mm; 21-May-2018, 1.5 m depth, 1 spc., adult, *L* = 45 mm; 1-Nov-2018, 1 m depth, 5 spcs, adults and egg masses, *L* = 15–30 mm; l’Espigó, Roses (Spain), 42°15’39.8”N, 3°10’26.8”E, 28-Feb-2020, 0.6 m depth, 8 spcs, adults and egg masses, *L* = 10–20 mm.

### External morphology

Body short, thick, background colour black. Mantle margin white with numerous appendixes black and yellow in colour, yellow close to apex, white papillae scattered all over body. Rhinophores lamellar, black forwards and white backwards, yellow on top. Gill leaves black, tips yellow.

### Ecology

All specimens found in each locality were found in shallow waters on the bryozoan *Bugula neritina* (Linnaeus, 1758), feeding and laying the egg masses on top.

### Distribution

Originally off the Pacific coast of America, this species is reported in Australia, Japan, South-Africa, Morocco (Moro *et al*., 2017); Cantabrian Sea (Cervera *et al*., 2004); and in the Mediterranean Sea in Italy, Croatia (Ballesteros *et al*., 2012–2020); France (GBIF, 2020; this study); Spain: Catalonia (this study).

### Remarks

Its prey grows on boat hulls, and thus, may be their way of propagation. Although this bryozoan is very abundant within Ports, *P. hedgpethi* has not been recorded in the biofouling samplings containing *B. neritina*. Some of the data on this species in the Mediterranean are in environments with variable salinity, such as the étang de Thau, or the Espigó de Roses, where there is freshwater supply.

> Genus *Polycerella* A. E. Verrill, 1880
>
> ***Polycerella emertoni* A. E. Verrill, 1880**
>
> (Figure 4G)

### Material examined

Cala Maset caves, Sant Feliu de Guíxols (Spain), 41°47’10”N, 3°2’44”E, 23-Aug-2017, 1 m depth, 11 spcs, juveniles, adults, and egg masses, *L* = 2–5 mm; Alfacs mussel farms, Sant Carles de la Ràpita, Montsià (Spain), 40°37’20.1”N, 0°39’48.5”E, 12-Sep-2018, 0.7 m depth, >20 spcs, adults and egg masses, *L* = 4–5 mm; le Ponton, étang de Thau, Sète (France), 43°25’28.5”N, 3°42’E, 1-Nov-2018, 1.2 m depth, 7 spc., adults and egg masses, *L* = 5–7 mm.

### External morphology

Body short, narrow, background colour white with irregular black and white yellowish punctuation. Gills composed of 3 leaves, with short, smooth papillae found behind. Rhinophores smooth.

### Ecology

All specimens found at each locality were found on the bryozoans *Amathia verticillata* where they lay the egg masses.

### Distribution

Invasive species originally from the east coast of North America (GBIF, 2020); now found in Morocco, Portugal, and the south of Spain (Cervera *et al*., 2004); France (this study) and Catalonia (Camps & Prado, 2018; this study).

### Remarks

Originally described from the North American Atlantic coast (Verrill, 1881), this species has been cited in much of the American Atlantic, and the Mediterranean Spanish and African coast. Its prey is commonly found attached to boat hulls. In the case of the specimens found in Catalonia, it has been recorded in ports or in bays and places with abundant nutrient accumulation.

> Genus *Thecacera* J. Fleming, 1828
>
> ***Thecacera pennigera* (Montagu, 1813)**
>
> (Figure 4H)

### Material examined

Cala Maset caves, Sant Feliu de Guíxols (Spain), 41°47’10”N, 3°2’44”E, 14-Feb-2018, 1.5 m depth, 1 spc., adult, *L* = 20 mm.

### External morphology

Body short, thick, background colour white with numerous black and yellow dots. Rhinophores lamellar, protected by two papillae, the posterior one longer than the frontal. Gill leaves discoid in shape, protected by long posterior papillae.

### Ecology

The singleton was found on top of an unidentified bryozoan species of the genus *Bugula*. Species with nocturnal activity. During the day the specimens are found hiding at the base of bryozoans.

### Distribution

Originally described from the North Sea (England; Montagu, 1813); Northeast American coast, Portugal and Italy (GBIF, 2020); Spain: Canary Islands, Cantabric Sea (Cervera *et al*., 2004), Catalonia (this study).

### Remarks

This species has a worldwide distribution and usually found in temperate waters (Dekker, 1986). In the Atlantic it has peaks of abundance where in a few days large numbers of specimens appear (Willan & Coleman, 1984).

> Family TRINCHESIIDAE F. Nordsieck, 1972
>
> Genus *Tenellia* A. Costa, 1866
>
> ***Tenellia adspersa* (Nordmann, 1845)**
>
> (Figure 5A)

**Figure 5.**
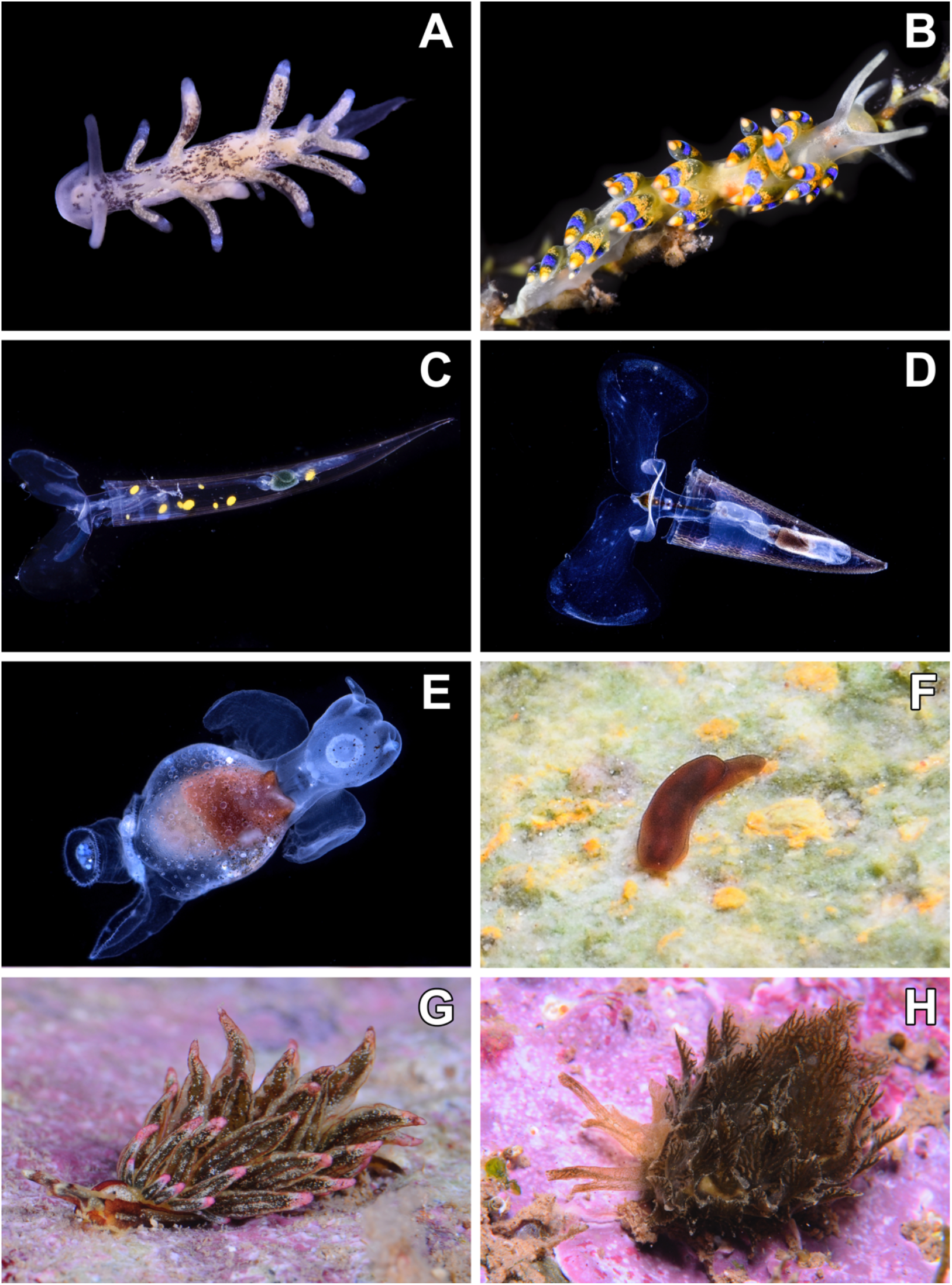
Live pictures of the Nudibranchia species: A *–Tenellia adspersa*. B – *Trinchesia cuanensis* on top of the hydrozoan *Sertularella* sp. Pteropoda: C – *Creseis virgula*. D – *Hyalocylis striata*. E – *Pneumodermopsis canephora*. Runcinida: F – *Runcina ornata*. Sacoglossa: G –*Aplysiopsis elegans*. H – *Caliphylla mediterranea*

### Material examined

Port de Blanes (Spain), 41°40’25.5”N, 2°47’48.6”E, 21-Jan-2019, 0.1 m depth, 1 spcs, adults, *L* = 7 mm.

### External morphology

Body elongate, narrow, background colour black. Rhinophores smooth, oral veil very developed, without oral tentacles. Cerata lateral, elongated; tip redounded.

### Ecology

The specimen was found in a mass of hydrozoans in a floating walkway with other heterobranchs.

### Distribution

Northeast Atlantic (OBIS, 2020); Pacific North American coast (OBIS, 2020); Portugal (Encarnação *et al*., 2020); Spain: Canary Islands, Atlantic Andalucian coast, Galicia, Levantine coast (Cervera *et al*., 2004), Catalonia (this study).

### Remarks

This species has a widespread and cosmopolitan distribution (Roginskaya, 1970), found in oceanic and brackish water areas (Thompson & Brown, 1984). This species is differentiated from other *Tenellia* species by having a characteristic oral veil connecting the oral tentacles and the cerata arranged in groups (Evertsen, *et al*., 2004). Normally the body and cerata are coloured in white but, depending on the diet, their colouration varies, as is the case in this study. Encarnação *et al*. (2020) found this species associated with the invasive hydrozoan *Cordylophora caspia* (Pallas, 1771) on artificial structures.

> Genus *Trinchesia* Er. Ihering, 1879
>
> ***Trinchesia cuanensis* Korshunova, Picton, Furfaro, Mariottini, Pontes, Prkić, Fletcher, Malmberg, Lundin & Martynov, 2019**
>
> (Figure 5B)

### Material examined

Punta del Romaní, L’Escala (Spain), 42°6’54”N, 3°10’9”E, 10-Feb-2014, 7 m depth, 1spec, juvenile, *L* = 3 mm; 14-Feb-2015, 14 m depth, 1spc., adult, *L* = 10 mm; Le Ponton, étang de Thau, Sète (France), 43°25’28.5”N, 3°42’E, 14-Apr-2018, 1 m depth, 3 spc., adults, *L* = 7–20 mm; Cala Maset caves, Sant Feliu de Guíxols (Spain), 41°47’10”N, 3°2’44”E, 15-Apr-2018, 0.6 m depth, 15 spcs, juveniles, adults, and egg masses, *L* = 4–15 mm; 6-Jun-2018, 0.6 m depth, 20 spcs, adults and egg masses, *L* = 10–15 mm; 11-Mar-2020, 0.5 m depth, 12 spcs, adults and mating, *L* = 7–15 mm; Tamariu beach (Spain), 41°55’0.5”N, 3°12’27.1”E, 11-Feb-2019, 2 m depth, 4 spcs, adults, *L* = 6–10 mm.

### External morphology

Body elongate, narrow, background colour white or light yellow. Rhinophores and oral tentacles smooth, more intensely white than the body. Red or orange mark between eyes and in central dorsum. Cerata globular, displaying a bright yellow-blue-yellow colouration.

### Ecology

Specimens were found mating and laying the eggs on top of the hydrozoans *Sertularella mediterranea* Hartlaub, 1901 and *Sertularella polyzonias* (Linnaeus, 1758) inside superficial caves.

### Distribution

Northeast Atlantic (Korshunova *et al*., 2019); Adriatic Sea (Ballesteros *et al*., 2012–2020); France (this study); Spain: Catalonia (this study).

### Remarks

This species was recently described and observed in the North Atlantic and the Mediterranean Sea (Korshunova *et al*., 2019). It is clearly different from other similar species of *Trinchesia* by having a red mark between the eyes and in the middle part of the back. It has a shorter life cycle than *T. morrowae* (Korshunova *et al*., 2019), since it has only been observed in the winter and spring months, whereas *T. morrowae* is observed all year round and *T. caerulea* (Montagu, 1804) only during spring and autumn and deeper (X. Salvador pers. obs.).

> Order PTEROPODA Cuviers, 1804
>
> Family CRESEIDAE Rampal, 1973
>
> Genus *Creseis* Rang, 1828
>
> ***Creseis virgula* (Rang, 1828)**
>
> (Figure 5C)

### Material examined

Ullastres 3, Llafranc (Spain), 41°53’14.1”N, 3°12’26.0”E, 27-Feb-2019, 0.5 m depth, 3 spcs, adults, *L* = 10 mm; Cala Sa Tuna, Begur (Spain), 41°57’38”N, 3°13’52”E, 9-Mar-2019, 0.8 m depth, 1 spec, adult, *L* = 10 mm; Montiel, Tamariu (Spain), 41°54’55.9”N, 3°12’46.3”E, 13-Mar-2019, 1 m depth, 20 spcs, adults, *L* = 5–10 mm.

### External morphology

Shell narrow, elongated, conical, back end slightly curved, smooth. Parapodia transparent, with the inner part thicker than the external.

### Ecology

Found in open waters, close to the surface around other pelagic pteropod and salp species.

### Distribution

Cosmopolitan, in the Mediterranean Sea this species is reported in the eastern part, and in the western, in the Malta stretch and between Mallorca and Sicilia (OBIS, 2020). This is the first report in the Mediterranean Spanish coast.

### Remarks

Almost all records in the Mediterranean are based on shell observations or are caught in plankton campaigns, in open waters and either at superficial depths (0–10 m) or at greater depths, (600–700 m; OBIS, 2020). Our specimens were found in February and March along the massive arrival of pelagic zooplankton at superficial waters.

> Genus *Hyalocylis* Fol, 1875
>
> ***Hyalocylis striata* (Rang, 1828)**
>
> (Figure 5D)

### Material examined

Cala Sa Tuna, Begur (Spain), 41°57’38”N, 3°13’52”E, 9-Feb-2019, 1 m depth, 1 spc, adult, *L* = 6 mm; Montiel, Tamariu (Spain), 41°54’55.9”N, 3°12’46.3”E, 13-Mar-2019, 1 m depth, 4 spcs, adults, *L* = 6– 10 mm; 14-Mar-2019, 1 m depth, 3 spcs, adults, *L* = 6–10 mm.

### External morphology

Shell wide, striated, conic, translucent. Parapodia large, transparent. Statocysts visible between shell and parapodia.

### Ecology

Found in open waters, close to the surface around other pelagic pteropod and salp species.

### Distribution

Cosmopolitan, in the Mediterranean Sea it is commonly reported in the eastern part (Koukouras, 2010; OBIS, 2020) and the Italian coast (OBIS, 2020). This is the first report in the Mediterranean Spanish coast.

### Remarks

As for *C. virgula*, almost all records in the Mediterranean are based on shells or specimens caught with plankton nets, in open waters and at great depths (OBIS, 2020). Our specimens were found in February and March along the massive arrival of pelagic zooplankton.

> Family PNEUMODERMATIDAE Latreille, 1825
>
> Genus *Pneumodermopsis* Keferstein, 1862
>
> ***Pneumodermopsis canephora* Pruvot-Fol, 1924**
>
> (Figure 5E)

### Material examined

Montiel, Tamariu (Spain), 41°54’55.9”N, 3°12’46.3”E, 13-Feb-2019, 1 m depth, 9 spcs, adults, *L* = 7–10 mm.

### External morphology

Body rounded, transparent, with numerous glands, dark chromatophores can be seen in by transparency all over body. Lateral gill extremely long, posterior gill absent. Posterior end presenting two ciliated bands. Head with 2 apical tentacles.

### Ecology

Found in open waters, close to the surface with other pteropods and proliferation of pelagic salps.

### Distribution

North Atlantic (OBIS, 2020); Mediterranean Sea (Pruvot-Fol, 1924); this the first report in the Catalan coast (Spain).

### Remarks

Mostly found in North Atlantic waters (Pruvot-Fol, 1924). Out of the eight specimens collected, two disappeared as well as some other species of pteropods collected inside the same container, likely meaning that in stressful situations they may be cannibalistic.

> Order RUNCINIDA
>
> Family RUNCINIDEA H. Adams & A. Adams, 1854
>
> Genus *Runcina* Forbes [in Forbes & Hanley], 1851
>
> ***Runcina ornata* (Quatrefages, 1844)**
>
> (Figure 5F)

### Material examined

Cala d’Aiguafreda, Begur (Spain), 41°57’49”N, 3°13’41”E, 13-Oct-2017, 0.1 m depth, 7 spcs, adults, *L* = 2–3 mm; 21-Nov-2017, 0.1 m depth, 1 spec, adult, *L* = 2 mm; 31-Jan-2018, 0.1 m depth, 9 spcs, adults, *L* = 2–3 mm; 23-Apr-2018, 0.1 m depth, 2 spcs, adults, *L* = 3 mm; 25-May-2018, 0.1 m depth, 2 spcs, adults, *L* = 3 mm; Punta del Romaní, L’Escala (Spain), 42°6’54”N. 3°10’9”E, 10-Jan-2018, 0.2 m depth, 4 spcs, adults, *L* = 2–3 mm; 24-Apr-2018, 0.2 m depth, 3 spcs, adults, *L* = 2–3 mm; 7-Dec-2018, 0.2 m depth, 7 spcs, adults, *L* = 2–3 mm; Punta d’en Bosch, Sant Feliu de Guíxols (Spain), 41°45’54”N, 3°0’11”E, 17-Apr-2018, 0.1 m depth, 5 spcs, adults, *L* = 1–3 mm.

### External morphology

Body elongate, narrow, translucid, black or brown in colour, rear tail darker. Foot large.

### Ecology

The specimens were found grazing on biofouling during the night over calcarean red algae or boulders, at very superficial waters.

### Distribution

Spain: Canary Islands and Strait of Gibraltar (Cervera *et al*., 2004), Catalonia (this study).

### Remarks

This species is rarely observed and very small. Easily distinguishable by the absence of spots and white marks and its uniform black colour (Quatrefages, 1844).

> Superorder SACOGLOSSA Ihering, 1876
>
> Family HERMAEIDAE H. Adams & A. Adams, 1854
>
> Genus *Aplysiopsis* Deshayes, 1853
>
> ***Aplysiopsis elegans* Deshayes, 1853**
>
> (Figure 5G)

### Material examined

Cova de l’infern (Spain), 42°19’2.75”N, 3°19’12.74”E, 19-Nov-2017, 1.2 m depth, 5 spcs, adults and egg masses, *L* = 15–20 mm; Cala Maset caves, Sant Feliu de Guíxols (Spain), 41°47’10”N, 3°2’44”E, 13-Sep-2019, 1.3 m depth, 1 spc., adult, *L* = 15 mm.

### External morphology

Body elongate, narrow, background colour white with longitudinal dark red lines in dorsum and laterals. Cerata long, smooth; background colour green whit white punctuation, dark red vertical lines, pink on tips.

### Ecology

The specimens were found feeding on the green alga *Cladophora prolifera* (Roth) Kützing, 1843. The egg mass is whitish, cylindrical, found inside the algal masses. The specimen observed in 2019 was found at night crawling over algae, while the 2017 specimens were hidden inside algae. This could mean that they are mostly nocturnal, hiding in the algal masses during the day.

### Distribution

Greece (OBIS, 2020); Croatia (Mavrič *et al*., 2014); Spain: Canary Islands (Ortea *et al*., 1998, 2001), Balearic Islands (Ballesteros & Templado, 1996), Catalonia (this study).

### Remarks

This species is rarely observed, found among algal samples, especially in *Cystoseira* and *Cladophora* as well as *Halopteris scoparia* (Linnaeus) Sauvageau, 1904 (Mavrič *et al*., 2014) and probably feeding on it.

> Genus *Caliphylla* A. Costa, 1867
>
> ***Caliphylla mediterranea* A. Costa, 1867**
>
> (Figure 5H)

### Material examined

Cala d’Aiguafreda, Begur (Spain), 41°57’49”N, 3°13’41”E, 5-Sep-2017, 0.8 m depth, 2 spcs, adults, *L* = 17 mm; CalaVentosa, Sant Feliu de Guíxols (Spain), 41°47’5”N, 3°2’52”E, 13-Sep-2017, 1.5 m depth, 1 spc., adult, *L* = 20 mm; Cala Maset caves, Sant Feliu de Guíxols (Spain), 41°47’10”N, 3°2’44”E, 5-Dec-2017, 0.7 m depth, 9 spcs, adults and egg masses, *L* = 5–40 mm; Fòrum pools, Barcelona (Spain), 41°24’34.5”N, 2°13’36.8”E, 5-Dec-2017, 0.7 m depth, 8 spcs, juveniles, adults, and egg masses, *L* = 5–40 mm; Cova de l’infern (Spain), 42°19’2.75”N, 3°19’12.74”E, 18-Sep-2018, 1.3 m depth, 1 spc., adult and egg masses, *L* = 40 mm.

### External morphology

Body elongate, thick, background colour light green with brown and white small dots along body, white pigmented area between eyes. Cerata numerous, flat; transparent with green lines running towards apex. Rhinophores long, folded, divided in “Y” shape, tips divided into two unequal parts.

### Ecology

The specimens have been found both at night and during the day on top of the green algae *Bryopsis duplex* (De Notaris, 1844) and *Cladophora prolifera* laying their egg masses. At night they were actively crawling and laying the egg masses, while by day they hide in the algal masses.

### Distribution

Eastern coast of North America; Brazil (GBIF, 2020); Gibraltar (García-Gómez, 2002); Spain: Canary islands (Moro *et al*., 2003; Ortea *et al*., 1999, 2001; Ortea & Bacallado, 1981), Andalucía (García Raso *et al*., 1992; Luque, 1983; Ocaña *et al*., 2000), Levantine coast (Templado *et al*., 1987, 2002), Balearic Islands (GROC, 2009–2020), Catalonia (this study).

### Remarks

This species is distinguished from other similar ones like *C. viridis* (Deshayes, 1857) (formerly *Polybranchia viridis*) by the presence of a ramified digestive gland in the cerata (see *Poybranchia viridis* (Rudman, 2020)).

> Genus *Cyerce* Bergh, 1870
>
> ***Cyerce graeca* Thompson T., 1988**
>
> (Figure 6A)

**Figure 6.**
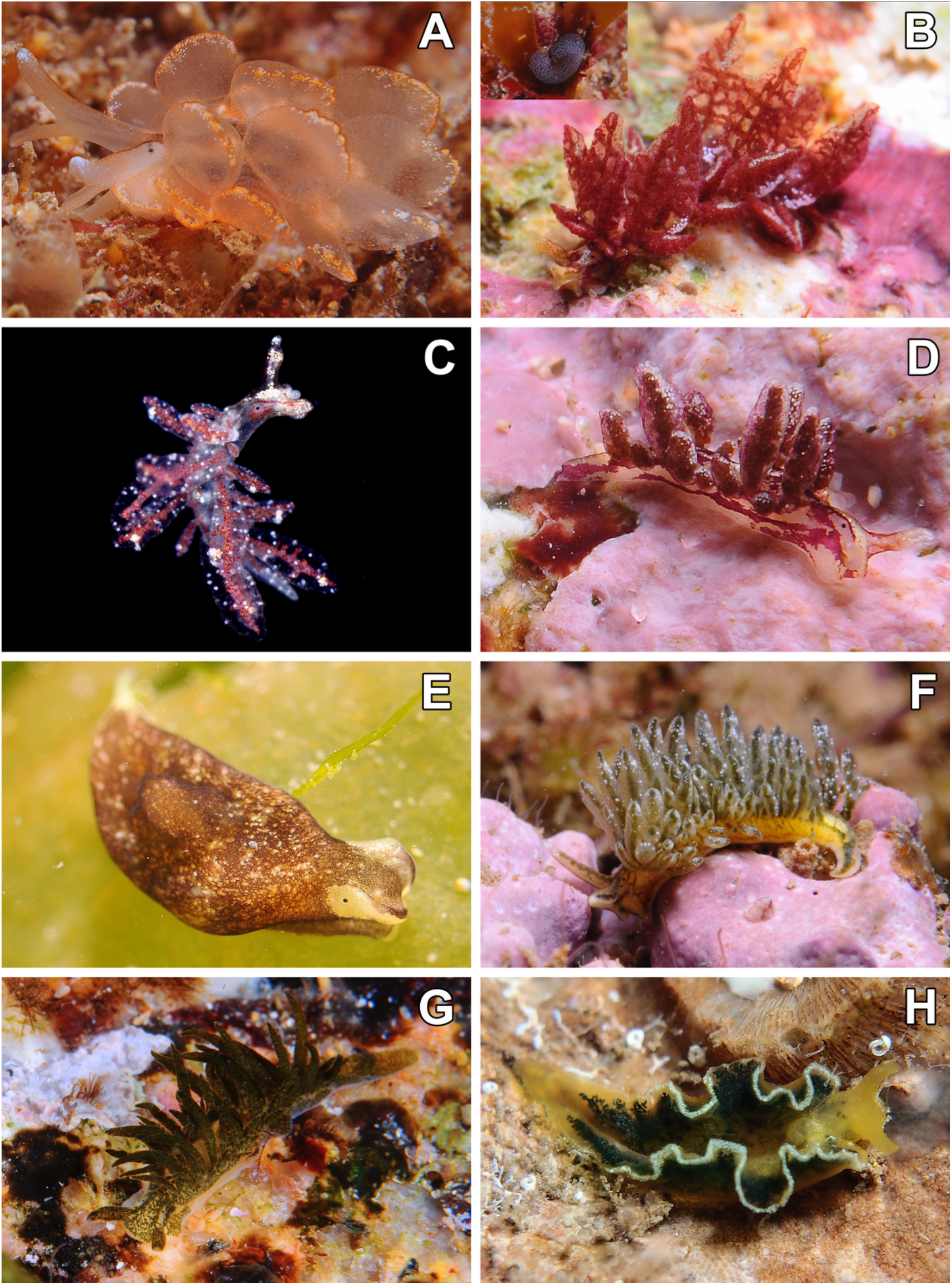
Live pictures of the Sacoglossa species: A – *Cyerce graeca*. B – *Hermaea bifida* and close-up on its egg mass at the base of an alga. C – *Hermaea cantabra*. D – *Hermaea paucicirra*. E – *Limapontia capitata* on *Ulva laetuca*. F – *Placida tardyi*. G – *Placida viridis*. H – *Elysia flava*.

### Material examined

Cala d’Aiguafreda, Begur (Spain), 41°57’49”N, 3°13’41”E, 26-Aug-2015, 0.5 m depth, 1spc., adult, *L* = 15 mm; 3-Aug-2017, 2.3 m depth, 1 spc., adult; Cala Ventosa, Sant Feliu de Guíxols (Spain), 41°47’5”N, 3°2’52”E, 13-Sep-2017, 1.3 m depth, 1 spc., adult; Cala Maset caves, Sant Feliu de Guíxols (Spain), 41°47’10”N, 3°2’44”E, 29-Jan-2018, 1.2 m depth, 1spc., adult; 14-Feb-2018, 0.7 m depth, 1 spc., juvenile, *L* = 3 mm; 22-Feb-2019, 1 m depth, 1 spec, adult, *L* = 12 mm; 20-Feb-2020, 0.7 m depth, 1 spec, adult, *L* = 10 mm.

### External morphology

Body elongate, thick, background colour transparent beige with darker notum. Cerata rounded, globular, beige, semi-transparent in colour, with blunt and dark brown digitations on top, white dots concentrated under these digitations. Rhinophores divided in “Y” shape.

### Ecology

All specimens were found on top of unidentified red algae; juveniles were found in February. Actively found crawling at night and usually under rocks in areas with abundant algae during the day.

### Distribution

Originally described from Greece (Thompson, 1988); Adriatic Sea and Italy (Ballesteros *et al*., 2012–2020); Spain: Balearic Islands (GROC, 2009–2020), Catalonia (Ballesteros *et al*., 2016; this study).

### Remarks

This species is more abundant in the eastern Mediterranean basin. Easily differentiated from other Mediterranean *Cyerce* species by the presence of a rounded parapodial margin and digitate cerata tips, whereas *C. cristallina* present an angular parapodial margin and roughened but not digitated cerata apexes (Thompson, 1988).

> Genus *Hermaea* Lovén, 1844
>
> ***Hermaea bifida* (Montagu, 1816)**
>
> (Figure 6B)

### Material examined

Cala d’Aiguafreda, Begur (Spain), 41°57’49”N, 3°13’41”E, 31-Mar-2017, 1 m depth, 9 spcs, adults, *L* = 7–12 mm; 25-Dec-2017, 1 m depth, 5 spcs, adults and juveniles; Cala Maset caves, Sant Feliu de Guíxols (Spain), 41°47’10”N, 3°2’44”E, 31-Dec-2017, 0.7 m depth, 2 spcs, juvenile and adult, *L* = 2–6 mm; 8-Mar-2018, 1.4 m depth, 2 spcs, adults and egg masses, *L* = 12 mm; Cala Sa Tuna, 41°57’38”N, 3°13’52”E, 3-Jan-2018, 0.8 m depth, 4 spcs, adults, *L* = 5–7 mm; Punta del Romaní, L’Escala (Spain), 42°6’54”N, 3°10’9”E, 6-Jan-2018, 0.7 m depth, 1spc., juvenile, *L* = 2 mm; Cala Trons, Lloret de Mar (Spain), 41°41’57”N, 2°51’52”E, 20-Jan-2018, 1.7 m depth, 1 spc., adult, *L* = 12 mm.

### External morphology

Body elongate, narrow, background colour red semi-transparent, with dispersed white dots more concentrated around head, cerata, and rhinophore apexes. Cerata globular, leave-shaped, semi-transparent, with internal lines of red colour running towards apex, this surrounded by four tubercles. Rhinophores beige, internally rolled, apex slightly longer than inferior part.

### Ecology

All specimens were found on top of the red alga *Bornetia secundiflora* (J.Agardh) Thuret, 1855 where they were mating and laying the eggs. The egg mass is whitish, cylindrical, and C-shaped (Fig. 5B, closeup); they were laid under the algae around, but never over *B. secundiflora*.

### Distribution

Central American coast and England (GBIF, 2020); Portugal (Cervera *et al*., 2004); Gibraltar (García-Gómez, 2002); Spain: Galicia (Rolán, 1983), Andalucía (Cervera *et al*., 1988), Levantine coast (Templado, 1982, 1984; Templado *et al*., 1983; Marín & Ros, 1988), Balearic Islands (GROC, 2009–2020), Catalonia (Ballesteros *et al*., 2016; this study); France (Rudman, 2020).

### Remarks

While in the Balearic Islands it was reported on the invasive alga *Lophocladia lallemandii* (Montagne) F. Schmitz, 1893 (GROC, 2009–2020), here we commonly found it on *B. secundiflora*. Juvenile specimens acquire a fluorescent orange colouration on the cerata, similar in colour to the latter alga.

> ***Hermaea cantabra* Caballer & Ortea, 2015**
>
> (Figure 6C)

### Material examined

Es Caials, Cadaqués (Spain), 42°17’5.1”N, 3°17’50”E, 20-Feb-2010, 10 m depth, 2 spcs, adults, H= 6–10 mm; Tamariu beach (Spain), 41°55’0.5”N, 3°12’27.1”E, 11-Feb-2019, 5 m depth, 1 spec, adult, *L* = 5 mm.

### External morphology

Body elongate, narrow, background colour semi-transparent, with two lateral and two dorsal red narrow lines close to eyes, diffuse at the end. Cerata globular, semi-transparent, with red digestive gland seen by transparency running towards apex, external white punctuations scattered but more concentrated into two tubercles before the top. Rhinophores beige in apical part, with a central red line and white dots scattered.

### Ecology

The specimen was found in a red filamentous algae sample at superficial depths.

### Distribution

North Spanish coast (Caballer & Ortea, 2015); Catalonia (GROC, 2009–2020; this study).

### Remarks

This species is distinguished from *H. bifida* by the presence of red lines that go from the rhinophores between the eyes to the back, the absence of oral appendages (present in *H. bifida*), and a less ramified digestive gland (Caballer & Ortea, 2015). The description of *H. cantabra* is recent thus, some previous records of *H. bifida* in the Iberian Peninsula could be misidentifications as for e.g., Ortea (1977) and Salvat (1968).

> ***Hermaea paucicirra* Pruvot-Fol, 1953**
>
> (Figure 6D)

### Material examined

Cala Maset caves, Sant Feliu de Guíxols (Spain), 41°47’10”N, 3°2’44”E, 5-Jan-2018, 0.4 m depth, 3 spcs, adults and copulation, *L* = 2–3 mm.

### External morphology

Body elongate, narrow, background colour translucent white, with red lines on dorsum going from apex of rhinophores to tail, and running on laterals from eyes, white opaque dots scattered on dorsum. Cerata club-shaped, red in colour, with white punctuation more concentrated in apex. Rhinophores internally rolled, back side longer than anterior part, transparent white whit opaque white dots.

### Ecology

The specimens were found feeding and mating on the red alga *Antithamnion cruciatum* (C.Agardh) Nägeli, 1847 and were active at night.

### Distribution

The description of this species occurred in Morocco (Pruvot-fol, 1953); Portugal (Cervera *et al*., 2004); Gibraltar (Cervera & García-Gómez, 1986); Spain: Canary Islands, Andalucía, Galicia, Cantabric Sea, Levantine coast (Cervera *et al*., 2004), Catalonia (Ballesteros *et al*., 2016; this study).

### Remarks

This species is distinguished from *H. bifida* by having a more globular and rounded cerata, and a more abundant white punctuation (Salvat, 1968). *Hermaea paucicirra* can be differentiated from its sympatric species *H. cantabra* by the presence of a more opaque white body, with a red mark on the eyes, and the because the red pigmentation on the dorsum barely allows to see the digestive gland as it also happen with the cerata (Caballer & Ortea, 2015).

> Family LIMAPONTIIDAE Gray, 1847
>
> Genus *Limapontia* Johnston, 1836
>
> ***Limapontia capitata* (O. F. Müller, 1774)**
>
> (Figure 6E)

### Material examined

Le Ponton, étang de Thau, Sète (France), 43°25’28.5”N, 3°42’E, 18-May-2017, 1 m depth, 2 spcs, adults, *L* = 3 mm.

### External morphology

Body short, thick, background colour dark brown, periocular area and tail white. Rhinophores very short, crest like. Parapodia absent. Protuberance caused by pericardial system seen in dorsum.

### Ecology

The specimens were found in brackish waters on bottoms extensively covered by *Ulva lactuca* Linnaeus, 1753.

### Distribution

Originally described is the North Sea (England and Norway; Muller, 1773); Portugal (Cervera *et al*., 2004); Spain: Cantabric Sea, Galicia, Levantine coast (Templado *et al*., 1983; Cervera *et al*., 2004); France (this study).

### Remarks

The species has been cited in other eutrophic areas with variable salinity, although it has been attributed to the fact that they were washed away by the waves (Rudman, 2020). It has always been found linked to green algae of the genus *Cladophora, Ulva*, and *Bryopsis*. It is distinguished from *L. senestra* (Quatrefages, 1844) by lacking long rhinophores and from *L. depressa* Alder & Hancock, 1862 by presenting this particularly shaped crests (Gascoigne, 1952).

> Genus *Placida* Trinchese, 1876
>
> ***Placida tardyi* (Trinchese, 1874)**
>
> (Figure 6F)

### Material examined

Fòrum pools, Barcelona (Spain), 41°24’34.5”N, 2°13’36.8”E, 23-Nov-2017, 0.4 m depth, 12 spcs, juveniles, adults, and egg masses, *L* = 3–20 mm; le Ponton, étang de Thau, Sète (France), 43°25’28.5”N, 3°42’E, 21-May-2018, 0.5 m depth, 3 spcs, adults, *L* = 17–25 mm; Cala Maset caves, Sant Feliu de Guíxols (Spain), 41°47’10”N, 3°2’44”E, 19-Dec-2018, 2 m depth, 2 spcs, adults and egg masses, *L* = 10–15 mm; 1-Jan-2019, 1.3 m depth, 3 spcs, adults, *L* = 13–15 mm; Cala Sa Tuna, Begur (Spain), 41°57’38”N, 3°13’52”E, 31-Jan-2019, 0.5 m depth, 1 spec, juvenile, *L* = 3 mm; Tamariu Beach (Spain), 41°55’0.5”N, 3°12’27.1”E, 10-Sep-2019, 1 m depth, 4 spec, adults with egg masses, *L* = 10–20 mm.

### External morphology

Body elongate, narrow, background colour white semi-transparent with a central white circle on back, laterals, and foot with dark brown or purple colour. Cerata long, smooth; background colour green, with white spots, tip dark garnet in colour. Rhinophores smooth, large, with white dots concentrated at the apexes. Oral tentacles wide and short.

### Ecology

The specimens were found in masses of the green alga *Bryopsis duplex*, mating and laying whitish and cylindrical egg masses at the base of the alga. This species appears completely unnoticed within the algal masses and moves actively during the night.

### Distribution

Originally described form Italy: Genoa (Trinchese, 1874), Gulf of Naples (Gascoigne & Sordi., 1980); Portugal (Calado *et al*., 2003); Gibraltar (Cervera *et al*., 1988); Spain: Catalonia (this study); France (this study).

### Remarks

This species can be misidentified with *P. vidiris* (Trinchese, 1874), but it is distinguished by the presence of brown or purple body margins and the tip of the cerata darkly coloured in garnet (Pruvot-Fol, 1954).

> ***Placida viridis* (Trinchese, 1874)**
>
> (Figure 6G)

### Material examined

Cala Ventosa, Sant Feliu de Guíxols (Spain), 41°47’5”N, 3°2’52”E, 13-Sep-2017, 0.1 m depth, 5 spcs, adults and egg masses, *L* = 7–20 mm; Cala Sa Tuna, 41°57’38”N, 3°13’52”E, 8-Mar-2018, 0.1 m depth, 6spcs, juveniles and adults, *L* = 2–15 mm; Punta d’en Bosch, Sant Feliu de Guíxols (Spain), 41°45’54”N, 3°0’11”E, 17-Apr-2018, 0.1 m depth, 5 spcs, adults and egg masses, *L* = 12–20 mm.

### External morphology

Body elongate, narrow, background colour white semi-transparent with digestive branches of dark green. Cerata long, smooth; background colour dark green, with white punctuation, apex white. Rhinophores long, translucent white with green branches and white punctuation on tips.

### Ecology

The specimens have been found on top of the green alga *Bryopsis mucosa* (J.V. Lamouroux, 1809) laying their cylindrical and white egg masses. Specimens in this study were found within a few centimetres’ depth feeding on *B. mucosa*, which is commonly found at subtidal depths.

### Distribution

Israel (Monselise & Mienis, 1977); Greece and Black Sea (OBIS, 2020); Italy (Ballesteros *et al*., 2012–2020); Spain: Catalonia (Cervera *et al*., 2004; this study); France (this study).

### Remarks

See Remarks section on *P. tardyi* above.

> Family PLAKOBRANCHIDAE Gray, 1840
>
> Genus *Elysia* Risso, 1818
>
> ***Elysia flava* Verrill, 1901**
>
> (Figure 6H)

### Material examined

Cala d’Aiguafreda, Begur (Spain), 41°57’49”N, 3°13’41”E, 30-Jul-2015, 2.3 m depth, 6 spcs, adults, *L* = 5–12 mm; Roqueo de los 14, La Herradura (Spain), 36°43’13.2”N 3°43’43.5”W, 18-Sep-2016, 1.6 m depth, 3 spcs, adults, *L* = 7–10 mm.

### External morphology

Body elongate, narrow, background colour green yellowish, dark green in parapodia, margin of parapodia wavy, white in colour. Rhinophores short, apex white, presenting two white spots between eyes.

### Ecology

The specimens were found mating in July on top of unidentified algae. This species is a strictly nocturnal, found during the day under rocks and at night on walls over masses of algae, especially green filamentous algae such as *Cladophora*.

### Distribution

Caribbean Sea (GBIF, 2020); Madeira (Cervera *et al*., 2004); Açores (Malaquias *et al*., 2009); Greece and Adriatic Sea (Ballesteros *et al*., 2012–2020); Spain: Canary Islands, Levantine coast, Catalonia (Cervera *et al*., 2004, this study), Andalucia (this study).

### Remarks

This species is clearly differentiated from other sympatric *Elysia* species by the yellowish colour of the body as well as the green dark colour in the parapodia (Thompson & Jaklin, 1988).

## Discussion

In the present study, we provide high-quality images, descriptions, and ecological remarks on 36 Mediterranean elusive species, corresponding to 20 new records in the Catalan coast (NE Spain). Our new data together with the recently described *Trinchesia morrowae* (Korshunova *et al*., 2019) and *Tylodina rafinesquii* (Fernández-Vilert *et al., in press*) increase the overall diversity of marine heterobranchs in this region up to 10%, with a total of 227 species. We also provide a new record from the southern Spanish Mediterranean coast, the sacoglossan *Elysia flava*. Moreover, the three species of pelagic pteropods are new records for Spain (i.e. *Creseis virgula, Hyalocylis striata* and *Pneumodermopsis canephora*) and 10 other species are new records for the French Mediterranean coast (i.e. *Haloa japonica, Haminoea orteai, Doto cervicenigra, Piseinotecus sphaeripherus, Polycerella emertoni, Amphorina andra, Trinchesia cuanensis, Limapontia capitata, Placida tardyi*, and *Placida viridis*). Regarding the alien species, we enlarge the knowledge on the distribution of five, *Haloa japonica* (in France), *Anteaeolidiella lurana* (in Spain: Catalonia and Valencia), *Godiva quadricolor* (in Catalonia), *Polycerella emertoni* (in France and Catalonia), and *Polycera hedgpethi* (in France and Catalonia).

Besides the new records of the native and invasive species, further information on the body morphology, phenology, egg mass morphology, and ecology of many species is provided. Briefly: regarding nudibranchs, numerous specimens of *Eubranchus prietoi* were firstly observed displaying lateralized cerata while being active. The specimen on *Aldisa smaragdina* reported here lacked the second dark circle present in its original description thus, suggesting a molecular revaluation of specimens of this genus would be useful to ascertain whether such chromatic variability corresponds to either population or species molecular diversity. Specimens of *Doto cervicenigra* were found along hydrozoans collected in harbours at one-meter depth. Another species of *Doto, D. pygmaea*, was found over floating objects (especially plastics) transported by sea currents and feeding on the hydrozoan *Tubelaria* sp., while only previously found feeding on *Obelia geniculata* (Linnaeus, 1758) by Ortea *et al*. (1997). Also, we found phenologic differences between the recently erected species of *Trinchesia*, while *T. cuanensis* have been observed in winter and spring, *T. morrowae* is present all year long, and *T*. *caerulea* only in spring and autumn. A rare species found in the Mediterranean, *Palio notus*, was regularly found in vertical walls that lacked direct sunlight, and it was found feeding on the bryozoan *Amathia lendigera*. Concerning the Pteropoda, the elusive *Pneumodermopsis canephora* was observed to have a cannibalistic behaviour, and life pictures of the three described species are some of the few published so far. Regarding sacoglossans, we provide new data on the diet of several species. For instance, specimens of *Hermaea bifida* were recorded here over the red alga *Bornetia secundiflora* for the first time (reviewed in Caballer & Ortea, 2015) at a few meters of depth, in vertical walls without sun radiation; when consumed, algal fronds acquired a fluorescent orange colour, also displayed in juvenile slugs.

Overall, we found 17 species that are active at night hours and most of the records were made in shallow depths while freediving, normally not going further down than 5 meters depth. Thereby, we believe many of the common records of heterobranch species living under rocks may actually have nocturnal activity, also that rare species may actually be common species with an opposite behaviour to our sampling methods (day hours, depths, and localities), and probably changing our habits would increase the diversity of a particular area, as we have seen in this study. Here we prove the importance of citizen science participation in this particularly attractive group, the heterobranchs, the diving community through the use of underwater photography and online platforms such as GROC (GROC, 2009–2020) or OPK (Ballesteros, *et al*., 2012–2020) represent a powerful tool to find new records, track the spread of alien species, and contributing to the increase the knowledge in ecology and phenology of this group. Up to date, specifically in our association (GROC, 2009–2020), a total of 194 collaborators have entered 23,660 observations, corresponding to 74,781 specimens belonging to 233 species for a total of 2,963 diving spots along the Spanish and southern French Mediterranean coast, way too far from the observations that our scientific community could have done alone.

## Acknowledgements

We are indebted to the nearly 200 citizen collaborators of GROC that regularly monitor the Catalan coast and adjacent waters, especially to Xavier Lindo for directing the association. Peter C. Kohnert is acknowledged for the taxonomic insight provided for the pteropod species. J. Moles postdoctoral fellowship was supported by the Alexander von Humboldt Foundation (Germany). This is the study #7 of the GROC Association.

